# Differential Maillard Sensitivity Of Exoproteins Favors Keratin Recovery During Sludge Biopolymer Extraction

**DOI:** 10.64898/2026.02.02.703216

**Authors:** Amrita Bhattacharya, Axel G Rosenvinge, Arsalen Esselami, Sandeep Rellegadla, Allan Omar Ghamlouch, Arya Van Alin, Johan Palmfeldt, Thomas Seviour

## Abstract

Proteins are an abundant extracellular polymeric substance (EPS) of activated sludge from wastewater. Hot alkalinization is implemented industrially to recover the EPS from sludge. We sought to assess the feasibility of protein recovery from alkali EPS. We detected a low abundance of bacterial proteins in the EPS. Human keratin was highly abundant and could also be recovered. Keratin was observed as a dominant and integral component of the activated sludge flocs, and it was thus not an extraction artifact. Alkali extraction promoted Maillard reaction between proteins and sugars and removed recoverable peptide signatures. Subsequent chemical modification, along with denaturation, impaired protein binding to ion exchange resins, making bacterial proteins inaccessible to isolation. Keratin has high resistance to Maillard reaction under extraction conditions and thus persists in the EPS. While Maillard modifies bacterial proteins, the resultant product, and possibly even keratin itself, are valuable recoverable byproducts from activated sludge.

**Synopsis:** The different sensitivities of proteins to Maillard reaction determines which proteins dominate alkaline extracellular polymeric substance (EPS) extract from activated sludge, with keratin dominating in alkaline EPS and an integral activated sludge component.

## 1. Introduction

Biological wastewater treatment produces large quantities of biowaste (i.e. sludge), which is costly to dispose of ^1^. However, wastewater sludge is rich in extracellular polymeric substances (EPS), i.e. proteins and polysaccharides, which have high intrinsic value^2^. Sludge EPS is potentially a renewable source of biopolymers and could help relieve challenges associated with current polymer use. ^3^ Accordingly, several demonstration-scale EPS recovery facilities have been installed to recover EPS from a subset of activated sludge called granular sludge (i.e. Nereda). In these installations the EPS is extracted at elevated pH and temperature (i.e. alkali EPS) and has properties like flame retardant, thickener and bio stimulant^456^that make it attractive for applications in building^7^ and agriculture^8^. The extracted alkali EPS constitutes a complex mixture of unidentified biopolymers, and this heterogeneity hampers its subsequent utilization. It is, however, highly enriched in proteins, with protein content ranging from 13.4±0.7 wt.% to 37.2±3.4 wt.% per mass volatile suspended solids ^9^. This is only slightly less than from recognized bacterial microbial protein (MP) systems aiming to produce proteins as feed for livestock and aquaculture ^10^. As demonstrated for cheese whey, a 70-fold increase in protein content can easily be achieved with downstream processing ^11^. Hence, protein recovery from sludge EPS is theoretically viable, in non-human contact applications, to elevate EPS in the value pyramid, e.g. for precision fermentation, foaming agents, foliar fertilizers ^12^. The challenge though is to achieve extracts, where the protein is concentrated and in a state that is amenable to downstream processing.

Furthermore, unlike whey, the proteome of alkaline extracted EPS is poorly understood, even though alkaline extraction is the EPS recovery process currently implemented at an industrial scale. Proteins are inherently labile and are highly susceptible to degradation, denaturation, and precipitation, especially at high pH and temperatures (i.e. 40□-80□C) ^13^. Only few proteins exhibit thermal or pH stability under the alkaline extraction conditions e.g. keratin ^14^. In conventional proteomics, protein integrity is thus preserved through a range of strategies (e.g. mild buffers, low temperature)^15 16^. Proteomic studies on activated sludge often do not differentiate between bulk and extracellular proteins as they often extract proteins using CEX(Cation Exchange chromatography) or TCA precipitation ^17,18^. Nonetheless, based on proteomics studies using these methods, bacterial membrane proteins and enzymes abundant in activated sludge ^18^.

In this study we aimed to characterize the alkaline EPS proteome and assess the viability of recovering extracellular proteins from alkaline EPS. A low abundance of cellular proteins was observed, and the most abundant protein was human protein keratin ^14^. Keratin was observed widely distributed throughout the activated sludge matrix and was the protein that could be efficiently isolated from the alkaline EPS. We thus demonstrated that high temperature alkalinization causes substantial bacterial protein loss through both Maillard reactions from activated sludge EPS, and protein degradation. Consequently, the EPS proteome becomes significantly compromised, rendering bacterial proteins inaccessible to recovery by conventional extraction and purification methods.

## 2. Materials and Methods

### 2.1 Alkaline Extraction

For thermal-assisted EPS recovery, the sample was incubated at 80 °C for 1 h in the presence of 0.1 M NaOH. The remaining steps (centrifugation, recovery of the EPS fraction) were performed as described in the standard protocol for EPS extraction ^1920^.

### 2.2 Alkaline Extraction (Cold Treatment)

An equal volume of 0.1 M NaOH was added to reach a final concentration of 0.05 M NaOH. Samples were incubated at room temperature (20–25 °C) for 3 h with gentle agitation (100–150 rpm) or alternatively at 4 °C for 12 h to minimize cell lysis. The mixture was centrifuged at 12,000 × g for 20 min at 4 °C, and the supernatant was collected as the alkaline-extracted EPS fraction.

#### Proteomics

The mass spectrometry proteomics data have been deposited to the ProteomeXchange Consortium via the PRIDE^21^ partner repository with the dataset identifier PXD073568.

\# Detailed methods are in the Supplementary Methods section.

## 3. Results and Discussion

### 3.1 Keratin dominates bacterial proteins across all seasons, and is not an artifact

Human proteins, specifically cytokeratin, dominated the alkaline EPS from both wastewater treatment plant sludges in summer and winter (Fig 1A), as determined by mass spectrometry based proteomics. Wastewater treatment plant 2 (WT2) sludge showed seasonally higher protein levels in extracted EPS than from wastewater treatment plant 1 (WT1) (see iBAQ scores in Fig S1). iBAQ (intensity-Based Absolute Quantification) score in quantitative proteomics estimates the absolute abundance of proteins from mass spectrometry data and it provides accurate comparison of protein amounts within a sample.

**Figure 1.**
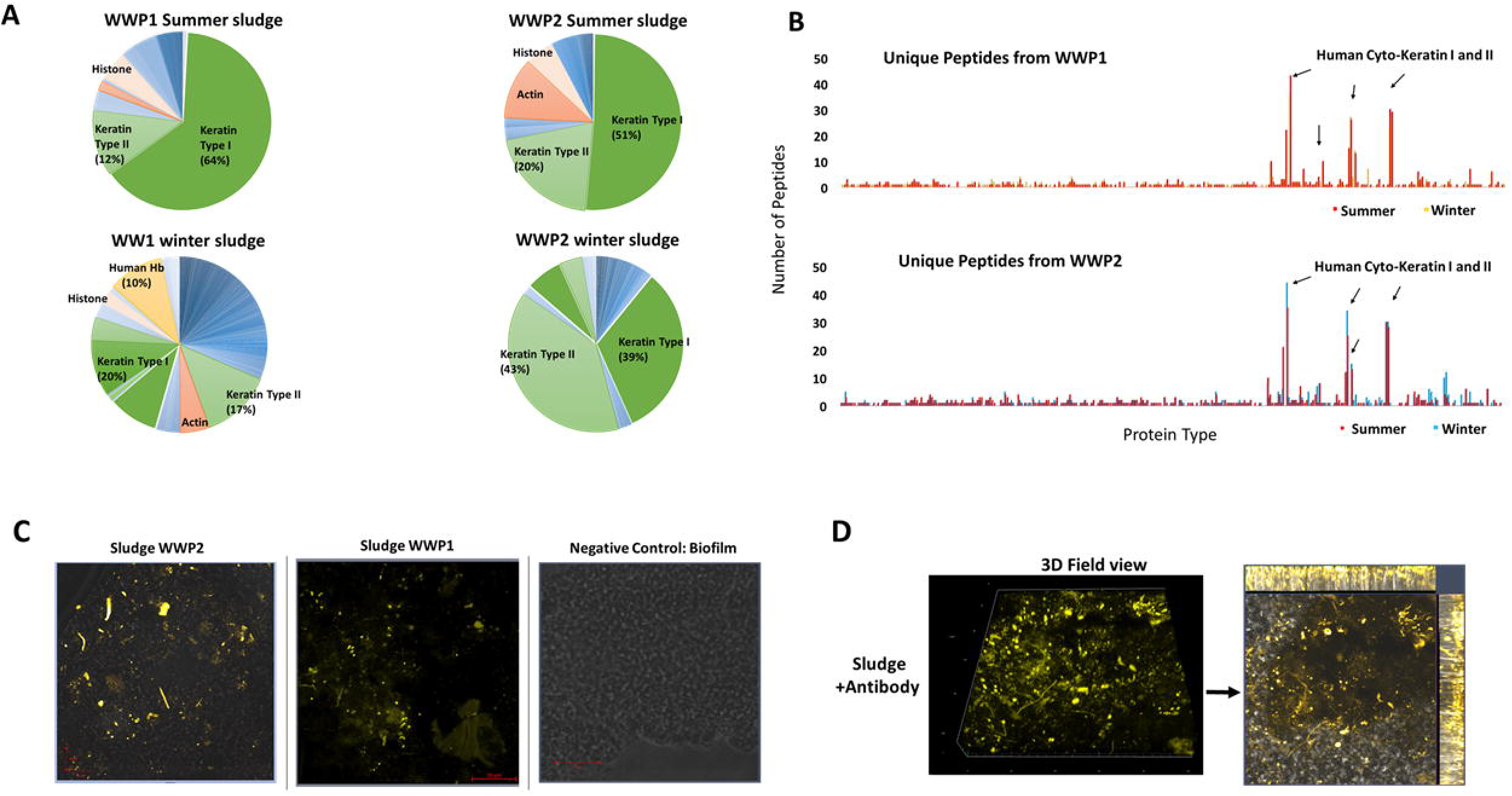
A) EPS protein percentages in alkaline EPS from WT1 and WT2 activated sludges (Percentage was calculated on all the iBAQ scores.) (B) Unique peptide scores of alkaline EPS from WT1 and WT2 activated sludges. (C) Confocal fluorescence micrographs of freeze-dried activated sludges from full-scale domestic Wastewater Treatment (WT) plants 1 and 2, and a suspended biofilm from a synthetic wastewater fed laboratory bioreactor (Negative control) stained with anti-keratin antibody specific for human cytoskeletal keratin (yellow) (n = 5). (D) Z-stack of confocal fluorescence micrographs of wet activated sludges from full-scale domestic wastewater Treatment (WT) plant 1, stained with anti-cytokeratin antibody. Each Z-stack image has 27 slices with each slice depth of 0.5μm.

Both sludges had more protein in winter samples than summer (Fig 1A), possibly due to reduced protein degradation. WT1 winter sludge had the most bacterial proteins. Additionally, there were more unique peptides for cytokeratin than for bacterial proteins for both WT1 and WT2 sludges (Fig 1B).

Immunofluorescence microscopy was performed directly on both summer sludges using human specific anti-cytokeratin antibody. Keratin was observed in all images collected and appeared distributed throughout the sludges (representative images shown in Fig 1C). It was not present in our control samples from biofilms from a bioreactor fed synthetic wastewater. Thus, the presence of keratin was not an artifact from the laboratory sample preparation. The dominance of keratin (and other human-derived proteins) in the wastewater EPS and its absence in the synthetic wastewater-fed bioreactor, suggests a strong anthropogenic contribution to the sludge extracellular protein pool.

Based on the z-stack of immuno-labelled sludge as shown in Fig 1D, keratin appeared integrated throughout the sludge matrix, thus indicating that it is assimilated into the flocs during bacterial aggregation Fig S2.

### 3.2 Alkaline extraction conditions promote Maillard reaction in EPS

As shown in Fig. 2A, a new molecular entity was created during sludge alkalinization. A high molecular weight (MW) peak appears in the exclusion chromatogram of sludge during alkalinization at 10.2 minutes retention time, corresponding to a MW of ≤50kDa. This occurred concomitant with a reduction in the intensity of the low MW peaks at 11.7 minutes retention time (i.e. <5kDa entity) (Fig 2A). Furthermore, the supernatant containing extracted EPS became increasingly brown over time (Fig 2B), corresponding to a UV-vis absorbance increase at ∼420 nm (Fig. 2C). This, as well as the formation of larger molecules from smaller molecules (Fig. 2A), is characteristic of the Maillard reaction, in which reducing sugars react with amino groups on proteins or peptides in sludge ^22^.

**Figure 2.**
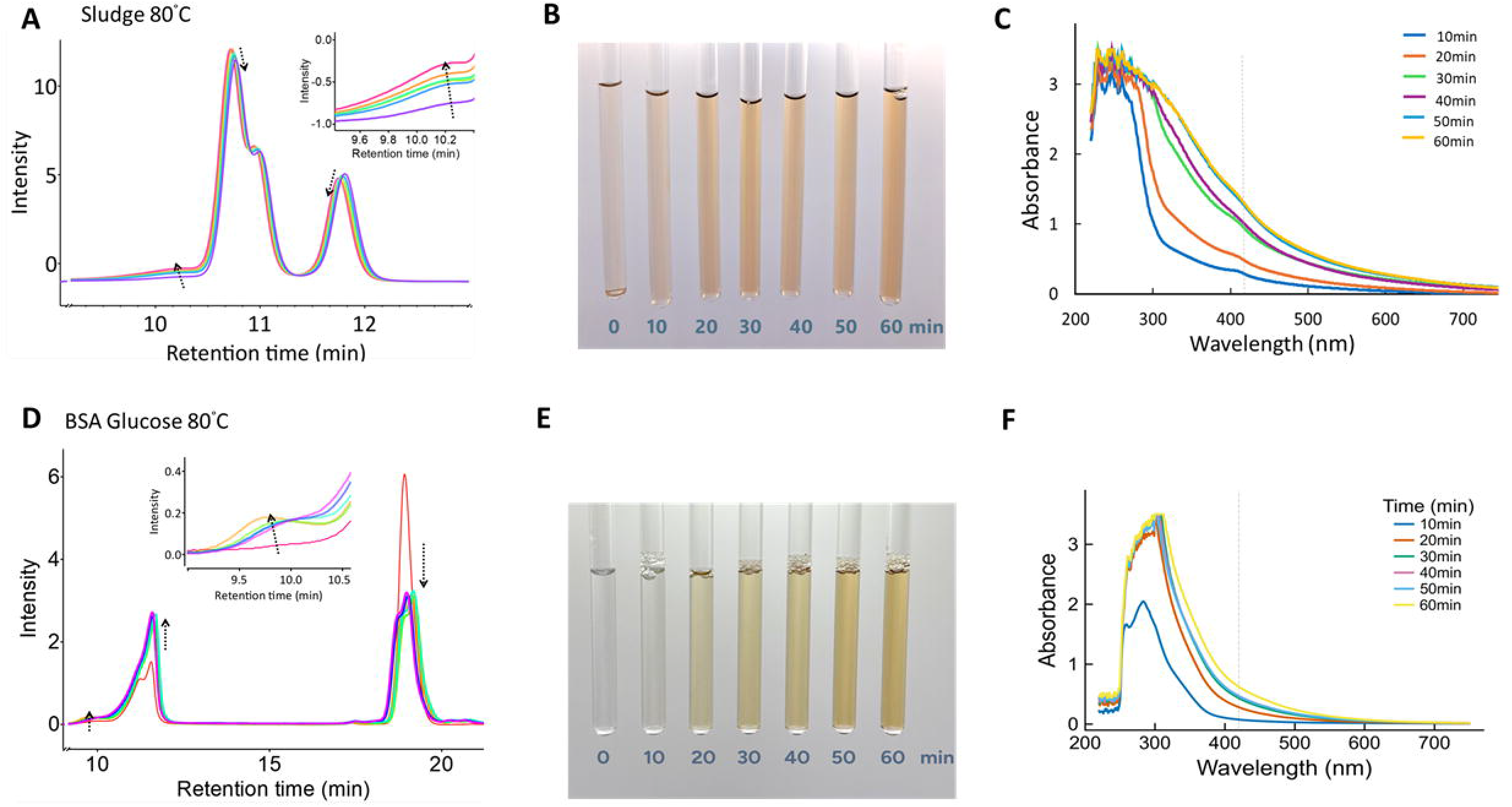
(A) High performance liquid chromatograms (HPLC) of WWT1 sludge supernatant upon sludge heating (pH 12, 0.1 mg sludge/mL) to 80 °C. Intensity is described in terms of RI signal (Arbitrary units). The insert shows highlights the intensity increase corresponding to the formation of a new high MW compound. (B) Sludge supernatant upon heating of activated sludge from WT1 under hot alkaline extraction conditions (T = 80 □ C) showing a color change. (C) UV-Visible range absorbance of alkaline EPS from WT1 sludge with wavelength scan range (220-750nm). (D) HPLC of bovine serum albumin (1mg/ml)/ glucose (1mg/ml) solution upon hot alkalinization (pH 12, T = 80 °C). The insert shows highlights the intensity increase corresponding to the formation of a new high MW compound. (E) Color change over time due to the reaction between BSA and Glucose under hot alkaline conditions. (F) UV-Visible range absorbance of the BSA-Glucose reaction.

We thus subjected a model system of BSA and glucose to the same extraction. A new high MW compound also formed (approximately 9.8 minutes retention time, size >100 kDa) (Fig 2D), similar to what was formed from sludge (inset Fig 2D). There was also a time-dependent increase in brown coloration (Fig. 2E), and increased absorbance from 300 – 600 nm. Together, these results indicate that the Maillard reaction occurs during sludge alkalinization and accounts for a loss of bacterial protein from EPS.

Maillard-driven glycation can enhance protein solubility and stability e.g. pea protein in food applications^23^. Although alkaline EPS extraction prevents bacterial protein isolation, it can nonetheless produce a more stable product still suitable for most EPS applications.

### 3.3 Keratin is the only protein that can be recovered from the alkaline EPS by cation exchange chromatography

As shown in Fig 3A, all proteins detected in the proteomic analysis of the sludges had isoelectric points between 4 and 11, with the majority falling within the 6-8 range. CEX can be used to isolate cationic proteins, which is the case for all proteins at pH < their isoelectric points. Thus, both acidic (bovine serum albumin, or BSA, pI 4.9) and basic (lysozyme, pI 11.0) proteins bound to the CEX column upon loading at pH 5 (as shown by an elution peak and the absence of a loading peak) (Fig S4).

**Figure 3.**
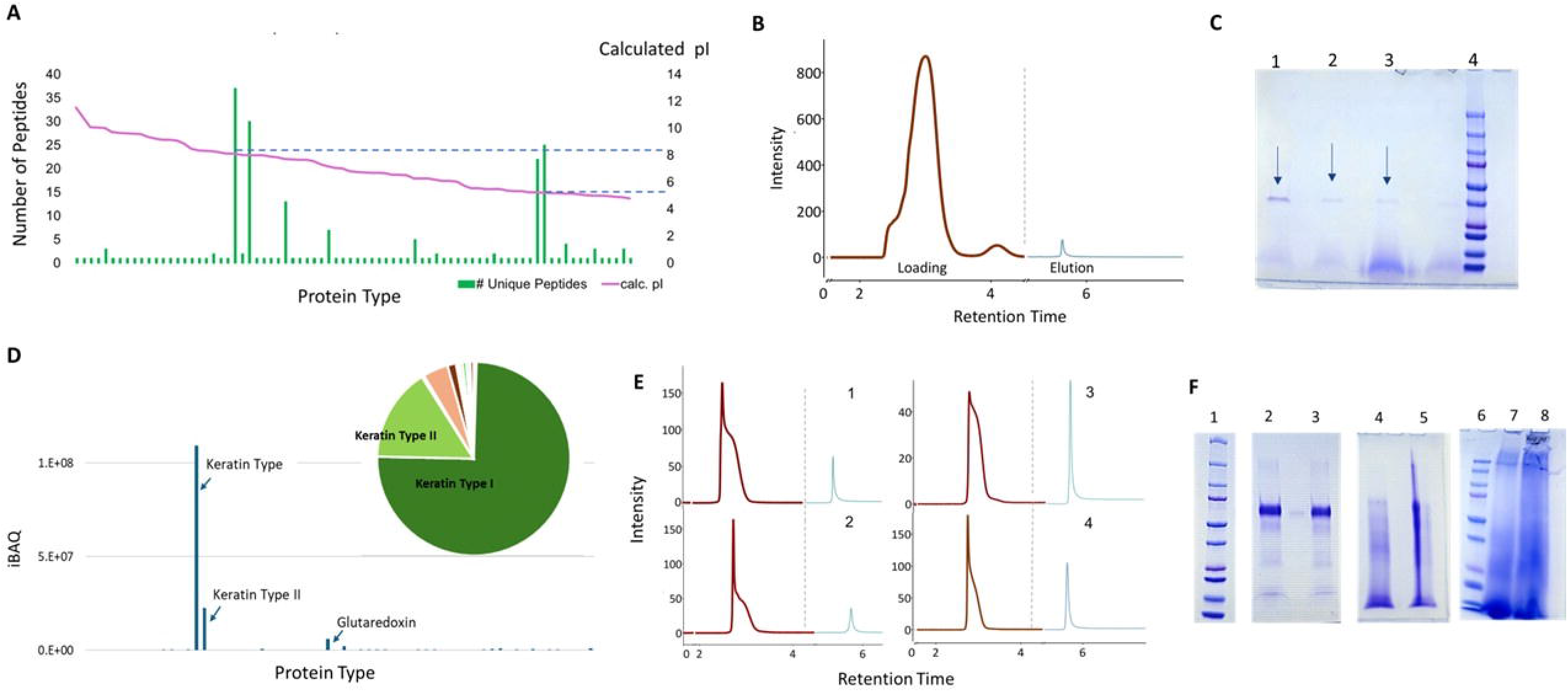
(A) The distribution of pI (Isoelectric point) of the peptides obtained from proteomic analysis of alkaline extracted sludge EPS. The purple line indicates that the pI range of the peptides is distributed from pH5 to pH 11. (B) Fast Protein Liquid Chromatography (FPLC) ultraviolet (UV) chromatogram of alkaline extracellular polymeric substances (EPS) extracted from sludge (Sludge with Hot Alkaline Extraction) after cation exchange chromatography (CEX). The wash fraction (Maroon line) represents unbound proteins, while the elution fraction (cyan line) represents proteins that were retained on and subsequently eluted from the CEX resin. (C) Sodium dodecyl sulfate–polyacrylamide gel electrophoresis (SDS–PAGE) of the concentrated elution fraction obtained from sludge HA reveals a clear protein band at ∼35 kDa.(Lane: 1)10X concentrated protein extract, 2) Protein Extract from wastewater plant WWP1, 3) Protein Extract from WWP2, 4) Ladder) (D) Quantitative proteomics (Q-proteomics) analysis of the excised SDS–PAGE band identified keratin as the predominant protein component, with only trace levels of glutaredoxin detected as shown by the iBAQ score. (E) Loading and elution profiles following cation exchange chromatography (CEX) showing the binding behavior of bovine serum albumin (BSA) to the CEX resin after hot alkaline (HA) and cold alkaline (CA) extraction. The figure also illustrates the extraction and subsequent CEX binding of a BSA–dextran mixture (1:1 ratio) following HA and CA extraction conditions. Plots: 1)BSA+Dextran HA, 2) BSA+Dextran CA, 3) BSA HA, 4) BSA CA (Loading(Maroon) and Elution(Cyan)) (F) Sodium dodecyl sulfate–polyacrylamide gel electrophoresis (SDS–PAGE) of bovine serum albumin (BSA) and a BSA–dextran (1:1) standard mixture subjected to hot alkaline (HA) and cold alkaline (CA) conditions, alongside alkaline-extracted sludge samples processed under the same HA and CA conditions. Lanes: 1) Ladder, 2) BSA CA, 3) BSA+Dextran CA, 4) BSA HA, 5) BSA+Dextran HA, 6) Ladder, 7) Sludge CA, 8) Sludge HA

The pH of the alkaline extract was reduced from 11 to 5 for loading the weak CEX column. No reduction in protein content was observed from the pH reduction (SI Fig S5). Nonetheless, only a small protein fraction was eluted from the alkaline EPS relative to the amount loaded under conditions suited to protein binding (i.e. pH 5) (Fig 3C). A clear band was obtained for the eluted protein on the SDS-PAGE gel and band intensity was proportional to concentration. The same band was oserved in the eluent after loading EPS from the WT2. A pure protein could thus be isolated from alkaline EPS despite the low elution signal (Fig 3C). Subsequent proteomic analysis of this protein revealed that the predominant component was keratin (Fig 3D). This is consistent with our previous finding that keratin dominated alkaline-extracted EPS.

CEX was then repeated under the same conditions (i.e. loading pH 5) on a model protein (BSA) pre-treated at pH 12 under conditions that do (80°C (HOT Alkaline or HA) and do not (T = 4 °C, Cold Alkaline or CA) promote Maillard, in the presence and absence of a reducing sugar. The presence of the reducing sugar reduced the amount of protein recovered by CEX under both conditions, but to a greater extent upon heating. Protein binding was reduced, however, under all alkali conditions, likely owing to protein denaturation. This loss of protein integrity was only observed upon heating (Fig. 3F). There were clear protein bands in the CA BSA sample with and without reducing sugar. HA BSA without reducing sugar appeared degraded. In the presence of the reducing sugar, however, a smear appears that extends above the MW of the BSA, indicating the formation of higher MW products as characteristic of Maillard. The same effects of smearing was also observed following hot alkali sludge treatment (Fig. 3F).

Finally, no color change in the keratin-glucose solution was observed upon hot alkaline conditions, indicating that the extraction conditions do not induce Maillard in Keratin (Fig S6(A) and Fig S6(B)). This further suggests that keratin has a high resistance to Maillard, likely explaining why it is recoverable from activated sludge alkali EPS.

Alkaline extraction of EPS from wastewater sludge selects for proteins with the greatest resistance to Maillard and denaturation, which in activated sludge is keratin. Alkaline conditions promote the Maillard reaction, hindering the recovery of labile proteins and creating fractions less favorable for conventional protein analysis and isolation. While alkaline extraction limits the isolation of the full proteome, it simultaneously transforms sludge EPS into a structurally resilient material, offering potential advantages for recovery and applications where stability and durability of the protein fraction are desirable

## Supporting information

Supplementary data and results

## Acknowledgement

This work was supported by funding from the Novo Nordisk Foundation under the ReThink Project, and WATEC. The authors sincerely thank these organizations for their support.

## References

[1] I. Karakas, S. B. Sam, E. Cetin, E. Dulekgurgen, and G. Yilmaz, “Resource recovery from an aerobic granular sludge process treating domestic wastewater,” Journal of Water Process Engineering, vol. 34, Apr. 2020, doi: 10.1016/j.jwpe.2020.101148.

[2] M. C. M. van Loosdrecht and D. Brdjanovic, “Anticipating the next century of wastewater treatment,” Science (1979)., vol. 344, no. 6191, pp. 1452–1453, Jun. 2014, doi: 10.1126/science.1255183.

[3] N. K. Kim, N. Mao, R. Lin, D. Bhattacharyya, M. C. M. van Loosdrecht, and Y. Lin, “Flame retardant property of flax fabrics coated by extracellular polymeric substances recovered from both activated sludge and aerobic granular sludge,” Water Res., vol. 170, p. 115344, Mar. 2020, doi: 10.1016/J.WATRES.2019.115344.

[4] T. M. Le, Y. Lin, W. Q. Zhuang, M. C. M. Van Loosdrecht, K. Jayaraman, and N. K. Kim, “Unlocking the potential flame-retardant mechanisms of extracellular polymeric substances recovered from municipal sludge,” J. Environ. Chem. Eng., vol. 13, no. 5, p. 117907, Oct. 2025, doi: 10.1016/J.JECE.2025.117907.

[5] Y. Tang, H. Xie, J. Sun, X. Li, Y. Zhang, and X. Dai, “Alkaline thermal hydrolysis of sewage sludge to produce high-quality liquid fertilizer rich in nitrogen-containing plant-growth-promoting nutrients and biostimulants,” Water Res., vol. 211, p. 118036, Mar. 2022, doi: 10.1016/J.WATRES.2021.118036.

[6] L. Świerczek, B. M. Cieślik, and P. Konieczka, “The potential of raw sewage sludge in construction industry – A review,” J. Clean. Prod., vol. 200, pp. 342–356, Nov. 2018, doi: 10.1016/J.JCLEPRO.2018.07.188.

[7] B. Pagliaccia, R. Campo, E. Carretti, M. Severi, C. Lubello, and T. Lotti, “Towards resource recovery-oriented solutions in agriculture exploiting structural extracellular polymeric substances (sEPS) extracted from aerobic granular sludge (AGS),” Chemical Engineering Journal, vol. 485, p. 149819, Apr. 2024, doi: 10.1016/J.CEJ.2024.149819.

[8] S. De Bruin, M. Riisgaard-Jensen, S. H. Hansen, M. C. M. Van Loosdrecht, P. H. Nielsen, and Y. Lin, “Global insights into extracellular polymeric substances from activated sludge: Yield, composition, and microbial communities,” Water Res., vol. 289, p. 124726, Jan. 2026, doi: 10.1016/J.WATRES.2025.124726.

[9] H. M. A. Shahzad, F. Almomani, A. Shahzad, K. A. Mahmoud, and K. Rasool, “Challenges and opportunities in biogas conversion to microbial protein: A pathway for sustainable resource recovery from organic waste,” Process Safety and Environmental Protection, vol. 185, pp. 644–659, May 2024, doi: 10.1016/j.psep.2024.03.055.

[10] C. Baldasso, T. C. Barros, and I. C. Tessaro, “Concentration and purification of whey proteins by ultrafiltration,” Desalination, vol. 278, no. 1–3, pp. 381–386, Sep. 2011, doi: 10.1016/j.desal.2011.05.055.

[11] Y. Yan et al., “Intracellular and extracellular sources, transformation process and resource recovery value of proteins extracted from wastewater treatment sludge via alkaline thermal hydrolysis and enzymatic hydrolysis,” Science of The Total Environment, vol. 852, p. 158512, Dec. 2022, doi: 10.1016/j.scitotenv.2022.158512.

[12] J. C. Bischof and X. He, “Thermal Stability of Proteins,” Ann. N. Y. Acad. Sci., vol. 1066, no. 1, pp. 12–33, Mar. 2006, doi: 10.1196/annals.1363.003.

[13] A. Shavandi, T. H. Silva, A. A. Bekhit, and A. E.-D. A. Bekhit, “Keratin: dissolution, extraction and biomedical application,” Biomater. Sci., vol. 5, no. 9, pp. 1699–1735, 2017, doi: 10.1039/C7BM00411G.

[14] A. V. Edhager, J. A. Povlsen, B. Løfgren, H. E. Bøtker, and J. Palmfeldt, “Proteomics of the Rat Myocardium during Development of Type 2 Diabetes Mellitus Reveals Progressive Alterations in Major Metabolic Pathways,” J. Proteome Res., vol. 17, no. 7, pp. 2521–2532, Jul. 2018, doi: 10.1021/acs.jproteome.8b00276.

[15] S. Fartade, T. Jadav, N. Rajput, and P. Sengupta, “A simplified optimization approach for sample preparation workflow in LC[MS[based quantitative proteomic analysis: Biological samples to peptides,” Arch. Pharm. (Weinheim)., vol. 358, no. 3, Mar. 2025, doi: 10.1002/ardp.202400911.

[16] P. Wilmes, M. Wexler, and P. L. Bond, “Metaproteomics Provides Functional Insight into Activated Sludge Wastewater Treatment,” PLoS One, vol. 3, no. 3, p. e1778, Mar. 2008, doi: 10.1371/journal.pone.0001778.

[17] P. Zhang et al., “Extracellular protein analysis of activated sludge and their functions in wastewater treatment plant by shotgun proteomics,” Sci. Rep., vol. 5, no. 1, p. 12041, Jul. 2015, doi: 10.1038/srep12041.

[18] M. Boleij, M. Pabst, T. R. Neu, M. C. M. van Loosdrecht, and Y. Lin, “Identification of Glycoproteins Isolated from Extracellular Polymeric Substances of Full-Scale Anammox Granular Sludge,” Environ. Sci. Technol., vol. 52, no. 22, pp. 13127–13135, Nov. 2018, doi: 10.1021/acs.est.8b03180.

[19] M. Boleij, M. Pabst, T. R. Neu, M. C. M. van Loosdrecht, and Y. Lin, “Identification of Glyoproteins Isolated from Extracellular Polymeric Substances of Full-Scale Anammox Granular Sludge,” Environ. Sci. Technol., vol. 52, no. 22, pp. 13127–13135, Nov. 2018, doi: 10.1021/acs.est.8b03180.

[20] Y. Perez-Riverol et al., “The PRIDE database at 20 years: 2025 update,” Nucleic Acids Res., vol. 53, no. D1, pp. D543–D553, Jan. 2025, doi: 10.1093/nar/gkae1011.

[21] N. Yang, S. Yang, and X. Zheng, “Inhibition of Maillard reaction during alkaline thermal hydrolysis of sludge,” Science of The Total Environment, vol. 814, p. 152497, Mar. 2022, doi: 10.1016/J.SCITOTENV.2021.152497.

[22] H. Khan, P. Mudgil, S. A. S. Alkaabi, Y. H. S. AlRashdi, and S. Maqsood, “Maillard reaction-based conjugation of pea protein and prebiotic (polydextrose): optimization, characterization, and functional properties enhancement,” Front. Sustain. Food Syst., vol. 8, Nov. 2024, doi: 10.3389/fsufs.2024.1463058.

