## Supplementary data and results for "Differential Maillard Sensitivity Of Exoproteins Favors Keratin Recovery During Sludge Biopolymer Extraction"

Supplementary Materials

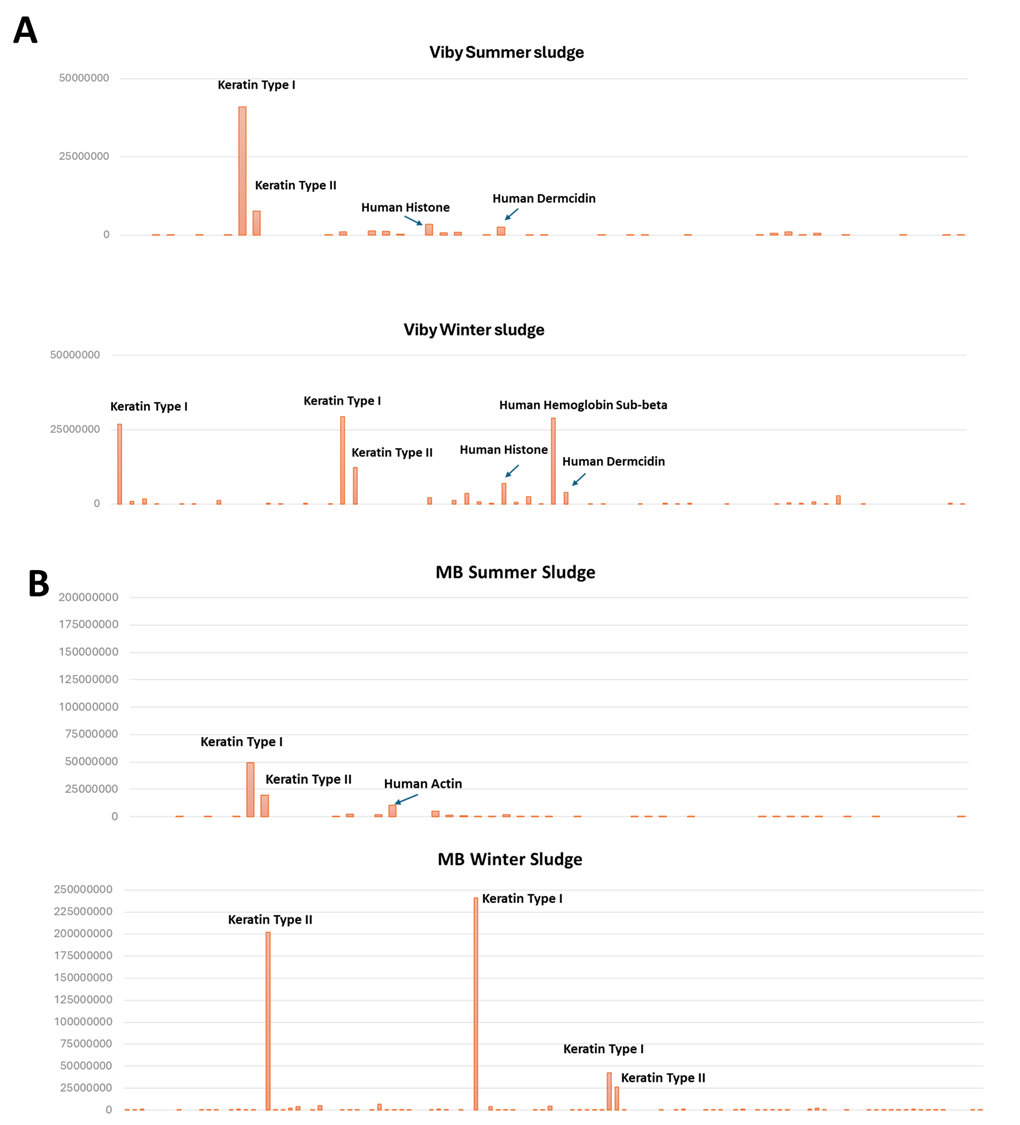

Fig S1: Comparison of protein in EPS from different sludges and seasons

A) Quantitative comparison using iBaq scores obtained from the Quantitative proteomics of Summer and Winter Sludge obtained from Viby

B) Quantitative comparison using iBaq scores obtained from the proteomics of Summer and Winter Sludge obtained from Marselisborg

Seasonally distinct sludge samples, collected during summer and winter from the Viby and Marselisborg wastewater treatment plants, were subjected to extracellular polymeric substance (EPS) extraction using alkaline-based methodologies. This approach enabled the isolation of EPS fractions, which are known to play a critical role in sludge structure, microbial community stability, and pollutant binding. The extracted EPS underwent downstream processing to remove residual sludge components other than proteins to prepare the samples for advanced molecular characterization using proteomic analysis.

The above graph represents the quantitative proteomcs data iBAQ score for each protein in different sludges across different seasons(Summer, Winter). Keratin family proteins are clearly more abundant than the other bacterial proteins.

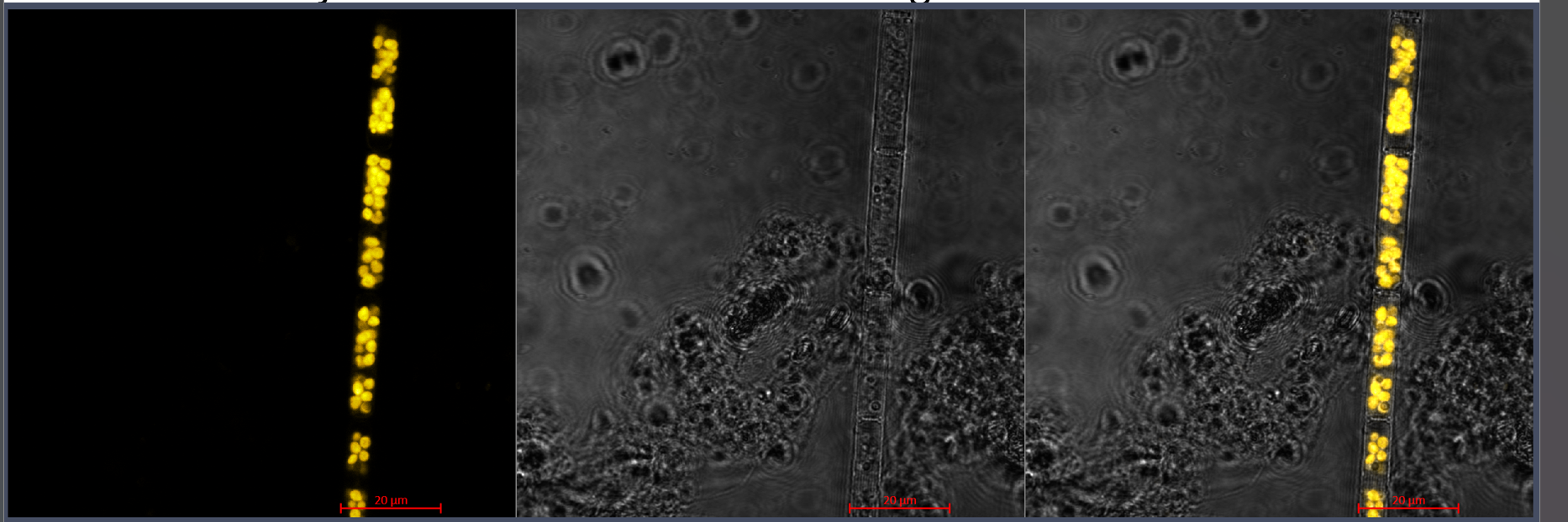

Figure S2: A) From left to right, the native autofluorescent signal of chlorophyll detected from a cyanobacteria, the brightfield image, and the merged image.

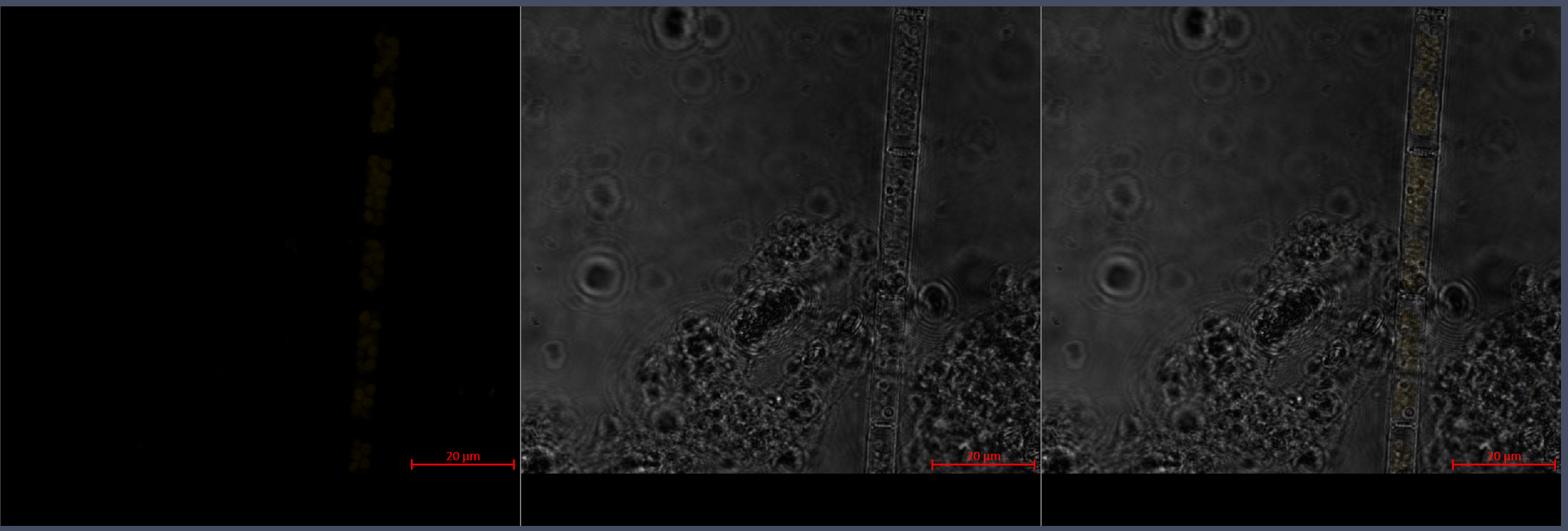

Figure S2: B) From left to right, the omitted autofluorescent signal (the detected signal is excluding any emission above 620 nm) of the cyanobacterial chlorophyll, the brightfield image, and the merged image.

Confocal laser scanning microscopy (CLSM) was performed using a Zeiss LSM700-AxioObserver system equipped with 20× EC Plan-Neofluar and 63× Plan-Apochromat oil immersion objectives (NA 1.4). Brightfield transmission imaging was used to visualise sample morphology. Fluorescence imaging was conducted using 555 nm excitation with peak emission at 585 nm for the keratin-specific tagged antibody. To minimise interference from chlorophyll autofluorescence of cyanobacteria or microalgae potentially present in the sludge, emissions above 620 nm were excluded during signal acquisition. Supplementary Figure S2 illustrates the effectiveness of this exclusion strategy, comparing the unfiltered native signal (Figure S2-B) with the adjusted signal (Figure S2-B).

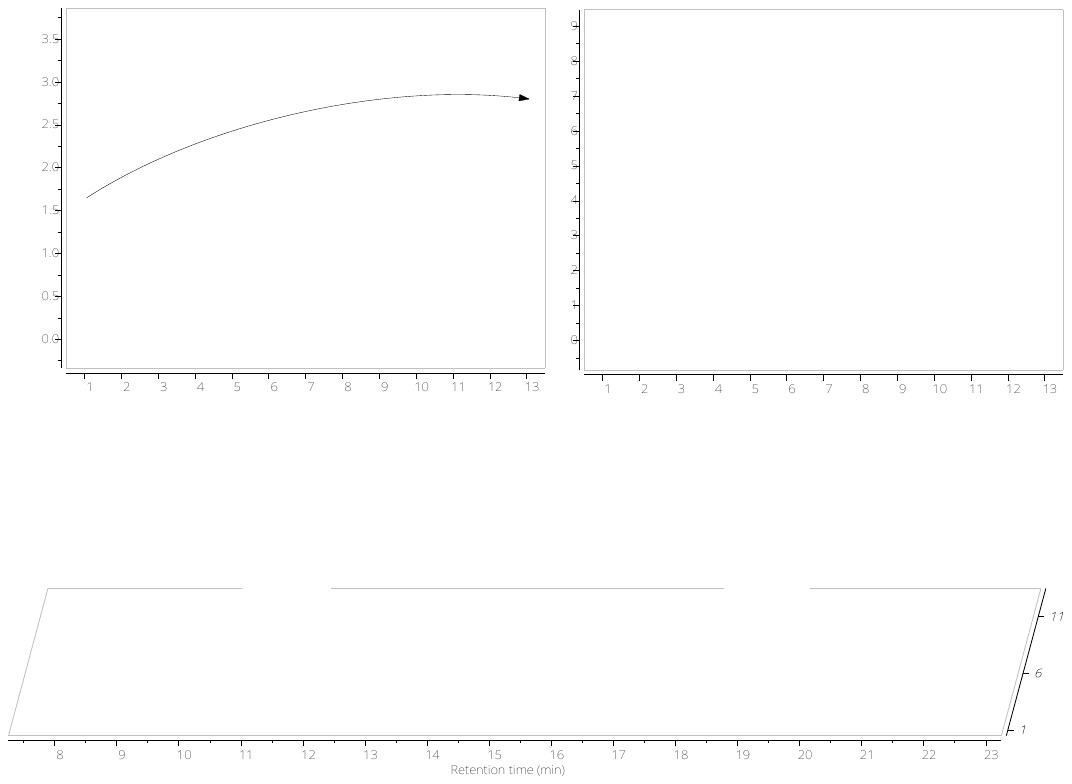

Fig S3: High-performance liquid chromatography (HPLC) was performed on alkaline extracts of BSA–glucose mixtures sampled at intervals of 5mins from 0 to 60 minutes. Here there is a a clear increase in intensity of one peak indicating formation of new product while there is a decrease in the other peak indicating glucose being utilized to modify BSA as the reaction progresses.

The HPLC analysis was performed using an OHpak SB-802.5 column (Shodex) to observe and separate smaller molecular weight products formed in the BSA–glucose system. This setup enabled the monitoring of molecular transformations occurring as a result of the Maillard reaction between BSA and glucose. The column’s resolution allowed effective detection of early-stage reaction intermediates and low-molecular-weight compounds, providing insights into the progression of product formation over time. Variations in peak intensities over the time intervals, along with the emergence of new peaks, suggest the formation of reaction products over time in the BSA–glucose system.

HPLC analysis of alkaline extracts of sludge samples was performed using an OHpak SB-806 column to observe and separate high–molecular-weight components. Over the various reaction timepoints between 0min to 60min, the appearance of a new peak indicated the formation or transformation of high–molecular-weight species within the sludge matrix over time during the alkaline extraction. This new species colud be due to the MRPs(Maillard reaction products) or intermediates which leads to an increase in brown colour over time.

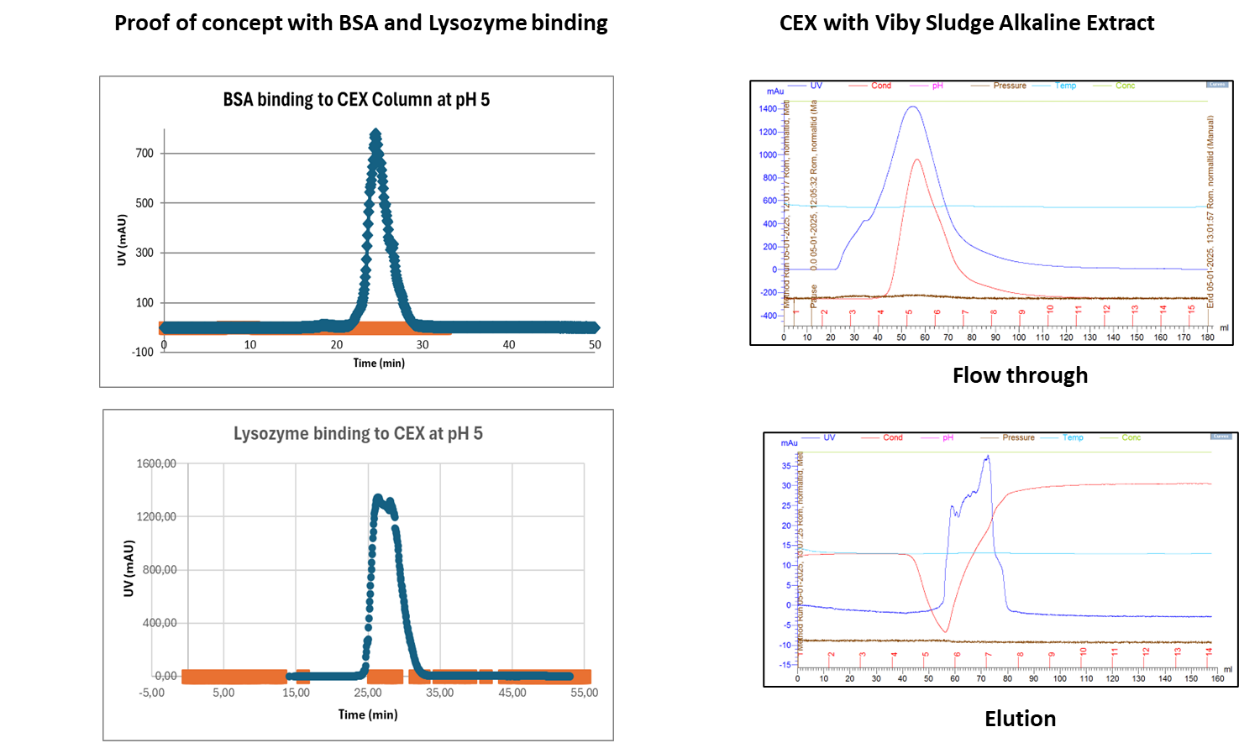

Fig S4: (A) Isolation optimisation of CEX(Cation exchange)resin at pH 5 using standard proteins Lysozyme and BSA.

(B) Isolation of proteins from Alkaine EPS extracted from Viby sludge: The blue line is the UV signal which in Flow Through indicates UV active molescules not binding to column and flowing out.

The Blue line is the UV signal which in Elution indicates binding of UV active components (which is then eluted).
