## Supplementary materials and methods for "Differential Maillard Sensitivity Of Exoproteins Favors Keratin Recovery During Sludge Biopolymer Extraction"

**Supplementary Methods**

**1. Protein Recovery and Precipitation**

For protein solubilization, the EPS extract (from any of the above methods) was diluted 1:1 with PBS (pH 7.4), and the pH was adjusted to 7.4 using 1 M HCl if necessary. Samples were mixed with 1% Triton X-100 and 10 mM DTT, gently shaken (100–150 rpm) at room temperature for 15–20 min, and sonicated in an ice bath using a probe sonicator (30–40% amplitude, 5s on/10 s off, total 3–5 min), ensuring the temperature remained below 30 °C. After sonication, centrifugation at 12,000 × g for 20 min at 4 °C yielded the supernatant containing solubilized proteins.

Proteins were purified by TCA–acetone precipitation. Trichloroacetic acid (TCA) was added to a final concentration of 10% (w/v), and the mixture was incubated at −20 °C for 30 min. The precipitated proteins were recovered by centrifugation at 12,000 × g for 20 min at 4 °C, and the pellet was washed with 1 mL of cold acetone, vortexed briefly, and centrifuged again at 12,000 × g for 10 min at 4 °C. The supernatant was discarded, and the pellet was air-dried at room temperature before being resuspended in Tris-HCl buffer (pH 6.8) containing 20 mM NaCl.

**2. SDS-PAGE Gel Electrophoresis**

**Sample Preparation**

The samples were mixed with a loading buffer (6X Laemmli SDS buffer) containing SDS, a reducing agent (like DTT or beta-mercaptoethanol), and a tracking dye (like bromophenol blue). 30ul of sample was added 6ul of loading dye to make the final concentration of 1X loading dye. The samples were then heated at 95^o^C for 10mins on a dry heating block. Then the samples were centrifuged at 8000 rpm for 5mins, and the supernatant was used to load the wells.

**SDS-PAGE:**

Precast SDS gels (Biorad) were used for analysis of the samples. The gel cassette was placed into the electrophoresis chamber (Biorad) and the chamber is filled with running buffer (1X-Tris-Glycine SDS buffer). An electric field of 100V was applied to run the gel. For the visualization of bands, the SDS-PAGE was run for around 1hour till the loading dye migrated to the bottom of the cassette. Then the gel was removed carefully from the cassette for staining. The protein bands were visualized after the staining and de-staining steps.

For quantitative proteomics the SDS-PAGE was only run to allow the sample to enter the stacking gel and then voltage was stopped. The gel was then removed carefully from the cassette, and the band was cut out using sterilized scalpel for proteomics analysis. This allowed us to obtain all the proteins and peptides together in a single band.

**SDS gel Visualization and Analysis:**

After electrophoresis, the gel was stained with Coomassie Blue stain for 1hour and then destained for 3-6 hours over gentle shaking conditions until clear bands were visible and the background stain was removed completely from the gels. The gels were then stored in distilled water overnight. The protein bands for proteomic analysis were cut, transferred to an Eppendorf tube, and stored at 4^o^C until the in-gel digestion procedure.

**Materials:**

SDS gel staining solution: The staining solution was prepared using Brilliant Blue R-250(Biorad) protein stain powder. The solution was prepared with 0.1% Coomassie Blue R-250, 40% methanol, and 10% acetic acid in distilled water.

SDS gel de-staining solution: The destaining solution recipe was prepared with 50% methanol, 10% acetic acid, and 40% water (v/v).

**3. Gel extraction of proteins for Proteomic Analysis**

i) In gel digestion from SDS-PAGE:

Label-free proteome preparations were essentially performed as previously described [(Edhager et al., 2018)]. The gel pieces were cut into 2X2mm pieces and were transferred to protein low bind Eppendorf tubes. They were then destained at 37^o^C for 30mins(twice), followed by reduction and alkylation of cystein residues. The gel pieces were shrunk using acetonitrile, and then trypsin (1µg/100 µL) in digestion buffer at pH 8. The sample was digested overnight at 37^o^C. The following day the supernatant was collected in a new Eppendorf (protein low bind) tube. Residual peptides from the gel pieces were extracted using acetonitrile and added to the supernatant. The supernatant containing the digested proteins was dried using a Speedvac and stored at -20^o^C.

ii) C-18 column purification of the proteins:

The lyophilized peptide/protein sample was dissolved in 100ul Sample buffer. The C-18 spin column (Thermo Scientific Pierce) was activated using activation buffer which was passed through twice and flow through was discarded post centrifugation (1500g for 1min). The whole sample was loaded on top of the resin bed of the C-18 column and column was centrifuged at 1500g for 1min. The flow through was discarded. The column was then washed twice with wash buffer and flow through was discarded post centrifugation (1500 g for 1 min). Sample was eluted from the C-18 column with elution buffer in Protein Lo Bind Eppendorf tube by centrifuge at 1500 x g for 1 min. The samples were lyophilized sample(s) (Dry the samples in the Speed Vac) and stored at -20^o^C until MS analysis.

**Materials:**

Reducing Buffer: Prepared just before use by mixing 10 µl of TCEP (500 mM) with 90 µl of Digestion Buffer for each digest to be performed. Final TCEP concentration is ~50 mM.(Freshly prepared)

Alkylation Buffer: Prepare just before use in foil-wrapped tubes to avoid exposure to light. Weigh 20 mg of Iodoacetamide (IAA) and dissolve in 200 µl water to make a 5X stock (~500 mM final concentration). Dilute the 200 µl of the 5X stock solution with 800 µl of Digestion Buffer to make the final Alkylation Buffer (50 mM). If more samples are being digested simultaneously, increase the volume of stock accordingly. (Freshly prepared)

Trypsin: Trypsin stocks were prepared from SIGMA / Promega. 100µg are dissolved in 100µl 50mM Acetic acid and portions of 10 ul stored in -20 freezer. Activated Trypsin was prepared Just before use dilute Stock trypsin solution 1:100 in digestion buffer and kept on ice.

Buffer for C18 purification**:** All buffers were prepared with MS grade Acetonitrile (ACN), water (H_2_O) and Methanol. Sample buffer composition was 5% ACN 0,5%TFA, Elution buffer was prepared with 70% ACN, MS H2O**.** Activation buffer was prepared with 50% MeOH, MS water.

**4. LC-MS/MS**

The peptide mixtures were analyzed by nano-liquid chromatography tandem mass spectrometry (nanoLC-MS/MS) (EASY nanoLC1200, Thermo Scientific) coupled to Q Exactive™ HF-X Quadrupole-Orbitrap™ Mass Spectrometer (Thermo Fisher Scientific)(Edhager et al., 2018) .The peptide samples were trapped on a pre-column (PepMap 100, 2 cm, 75 μm i.d., 3 μm C18 particles, 100 Å, Thermo Scientific) followed by reverse phase separation on a C18 column with integrated emitter (EASY-Spray column, PepMap 25 cm, 75 μm i.d., 2 μm, 100 Å, Thermo Scientific). Peptides were separated in a 60 min linear gradient, from 4 to 40% acetonitrile in 0.1% formic acid at a flowrate of 300 nL/min. The MS was operated in positive, data dependent mode, automatically switching between precursor scanning (MS1) and fragmentation (MS2) acquisition. Resolution of MS1 was set to 60,000 and MS2 to 45,000. In MS1, automatic gain control target was set to 1 × 10^6^ ions and scan range between 372 and 1500 m/z. In MS2, AGC target was set at 2 × 10^5^ ions, with fixed first mass set to 120 m/z. Dynamic exclusion was set to 20 s for all analyses. Up to twelve of the most intense ions were fragmented per full MS scan, by higher-energy C-trap dissociation. Ions with single charge or unassigned charge states were excluded from fragmentation.

**5. LC-MS/MS proteomics database searches**

Proteins were identified and quantified using MaxQuant^70^ (version 1.5.3.30) with its Andromeda algorithm against a sequence database of the following species: *Escherichia coli* (4589 entries downloaded 09/20/2023), *Homo sapiens* (20,398 entries, 08/22/2022), *Candidatus accumulibacter phosphatis* (17,061 entries, 02/01/2024), *Candidatus Phosphoribacter hodrii* (6,929 entries, 02/01/2024), *Pseudomonas aeruginosa* (3171 entries, 02/01/2024), *Zoogloea* species (29,566 entries, 02/01/2024). The “match between runs” was applied and iBAQ data were used for quantification.

**6. Immunolabelling and Confocal Microscopy of Keratin**

Keratin detection in the samples was carried out using a fluorescence-tagged specific antibody. A mouse IgG1 Pan Cytokeratin Monoclonal Antibody (AE1/AE3) tagged with eFluor™ 570 (0.2 mg/ml in phosphate-buffered saline (PBS), ThermoFisher) was used which can recognise many of the acidic and basic cytokeratin family members(Tahkola et al., 2021). For immunolabelling, the antibody was diluted in PBS containing 3% bovine serum albumin (BSA), which served as a blocking agent to minimise non-specific binding.

Pure keratin was used as a positive control, while samples from anammox biofilm served as negative controls. Both control and activated sludge samples were pre-incubated in 3% BSA in PBS for 2 minutes before the addition of the antibody (final working dilution 1:100 in 3% BSA in PBS). Samples were then incubated at room temperature with gentle agitation (50 rpm) for 60 minutes. Following incubation, the samples were rinsed with 3% BSA.

Confocal laser scanning microscopy (CLSM) was performed using a Zeiss LSM700-AxioObserver system equipped with 20× EC Plan-Neofluar and 63× Plan-Apochromat oil immersion objectives (NA 1.4). Brightfield transmission imaging was used to visualise sample morphology. Fluorescence imaging was conducted using 555 nm excitation with peak emission at 585 nm for the keratin-specific tagged antibody. To minimise interference from chlorophyll autofluorescence of cyanobacteria or microalgae potentially present in the sludge, emissions above 620 nm were excluded during signal acquisition. Supplementary Figure 2 illustrates the effectiveness of this exclusion strategy, comparing the unfiltered native signal (Figure S2-a) with the adjusted signal (Figure S2-B).

**7. Maillard Reaction**

### Preparation of the Glycine–Glucose Maillard Reaction Assay

To investigate the potential occurrence of the Maillard reaction and the effect of temperature and incubation time, BSA and glucose were mixed in a 1:2 molar ratio in potassium phosphate buffer at pH 12. The final concentrations in the reaction mixture were $3.37\cdot{10}^{-5}$ M and $5.99\cdot{10}^{-2}$ M, for BSA and glucose respectively with a total reaction volume of 5 mL. All samples were prepared in Eppendorf tubes to ensure consistent handling and minimize evaporation. Samples were incubated at 80 °C for 1 hour.

**Sample Preparation for Sludge**

To observe changes due to Maillard during Alkaline extraction 1mg dry sludge was added to 1ml pH12 Potassium phosphate buffer solution in 2ml Eppendorf tubes. The tubes were vortexed to mix sludge and the pH12 solution homogeneously and then the tubes were placed on a heating block at 80^o^C for 0min, 10min, 20min, 30min, 40min, 50min and 60min respectively. The Eppendorf tubes were removed at the said timepoints and centrifuged for 3mins at 10000rpm. The supernatant was removed to a fresh Eppendorf from each tube, and the colour change was observed visually.

### UV-Vis

UV–Vis absorbance spectra were recorded using a Merck Millipore Spectroquant Prove 600 spectrophotometer equipped with a monochromator. PMMA cuvettes (minimum filling volume 1.5 mL; Sigma-Aldrich) were used for all measurements. Spectra were collected over the wavelength range of 220–750 nm with a scan resolution of 1 nm. Samples were measured at room temperature, and a blank was recorded for baseline correction.

### Size exclusion chromatography for Glycine Glucose

High-performance liquid chromatography (HPLC) analyses were performed using a Waters e2695 separation system equipped with an OHpak SB-802.5 HQ column (300 × 8 mm). Milli-Q water served as the mobile phase at a flow rate of 0.6 mL min⁻¹ and a column temperature of 30 °C. Detection was carried out using both a Waters 2414 refractive index (RI) detector and a Waters 2489 UV detector. All samples were filtered through 0.45 µm syringe filters prior to injection. Chromatographic data was processed and analyzed using MestReNova 16 with the EIVis plugin.

### Size exclusion chromatography for Sludge

High-performance liquid chromatography (HPLC) analyses were performed using a Waters e2695 separation system equipped with an OHpak SB-806 HQ column. NaOH solution in Milli-Q water (pH 12) served as the mobile phase at a flow rate of 0.6 mL min⁻¹ and a column temperature of 30 °C. Detection was carried out using both a Waters 2414 refractive index (RI) detector and a Waters 2489 UV detector. All samples were filtered through 0.45 µm syringe filters prior to injection. Chromatographic data was processed and analyzed using MestReNova 16 with the EIVis plugin.

**Materials:** The reagents used in this study, potassium hydroxide (pellets), potassium phosphate stock solution (500 mM, pH 8), bovine serum albumin (BSA), and glucose were acquired from commercial suppliers. Potassium hydroxide (pellets) was purchased from Fisher Scientific and used to prepare a 0.25 M base stock for pH adjustment. A potassium phosphate stock solution (500 mM, pH 8) was obtained from Cayman Chemical Company, BSA was purchased from Sigma-Aldrich, and glucose was purchased from VWR Chemicals. All solvents were reagent graded, and all chemicals were employed as received from commercial sources

**8. Cation Exchange Chromatography**

Cation exchange chromatography (CEX) was performed using an ÄKTA Pure Fast Protein Liquid Chromatography (FPLC) system (Cytiva) equipped with a 40 mL column packed with SP Sepharose™ Fast Flow strong cation exchange resin and a UV detector operating at 280 nm. Prior to sample loading, the column was conditioned with three column volumes (CV) of 50 mM sodium acetate buffer (50 mM, pH 5). Subsequently, 25 mL of the conditioned sample was loaded onto the column. The column was then washed with three CV of 50 mM sodium acetate buffer (50 mM, pH 5) to remove unbound material. The tubes were flushed of excess sample before eluting three CV of 100 mM glycine buffer (100 mM, pH 12). Following elution, the column was cleaned with four CV of 2 M sodium chloride and finally washed with five CV of Milli-Q water to remove residual salts.

**Sample Preparation for FPLC:**

Extracellular polymeric substances (EPS) were extracted using an alkaline treatment procedure adapted from established protocols (Boleij et al., 2018). Dry sludge was dispersed in sodium carbonate solution (pH 11.5) and mixed by vortexing for 30 seconds, yielding a sludge concentration of 0.01 $g*mL^{-1}$.

Samples were subjected to one of two treatment conditions: incubation in a water bath at 80 °C for 1 hour (hot alkaline extraction) or stored at 4 °C overnight (cold alkaline extraction). Following incubation, the samples were centrifuged at 12,000 rpm for 20 minutes at room temperature. The supernatant was carefully collected, and the pellet was discarded. Sodium acetate (3 M, pH 5) was then added to the supernatant to adjust the pH to pH5(approximately). The solution was filtered through a 0.45 µm syringe filter to remove remaining particulates.

The standards were subjected to the same extraction protocol as the sludge samples. The standards consisted of bovine serum albumin (BSA) and a BSA–dextran mixture. The BSA extraction solution had a concentration of 1 $mg*mL^{-1}$.while the BSA–dextran solution contained the same overall protein concentration with a BSA:Dextran ratio of 1:3.

Determination protein content of pH reduced samples

The extraction solution was adjusted to a series of target pH values by the gradual addition of 3 M sodium acetate (pH 5) while continuously monitoring the pH with a calibrated pH meter. At each desired pH interval, a 500 µL sample was collected for analysis. Protein concentrations in all samples were determined using the Qubit Protein Quantification Assay, following the manufacturer’s instructions for calibration and standard curve generation.

The protein content in the solution was expressed as mass (mg) and calculated from the concentration determined using the Qubit assay by multiplying the measured protein concentration (mg/mL) by the sample volume (mL).

$$Mass of protein\left( mg \right)=concentraion of protein \left( \frac{mg}{mL} \right)*volume of sample\left( mL \right)$$

**Materials:**

Buffer solutions for the CEX experiments were prepared from sodium hydroxide (NaOH) and sodium chloride (NaCl), Merck. Glycine and sodium acetate were obtained from Sigma-Aldrich. An HQ11D portable pH meter was used to adjust the buffer pH. Captiva syringe filters from Agilent, compatible with various syringe volumes were used for filtration of samples. Sodium carbonate decahydrate (VWR), dextran (TCI), and bovine serum albumin BSA (Sigma Aldrich) used for sample preparation during alkaline extraction. Additional reagents were sourced from Sigma-Aldrich. The activated sludge was kindly provided by Aarhus Vand’s wastewater treatment plants in Viby and Marselisborg. Heating for the alkaline extraction process was in a Heto Shaking SBD50 water bath from Holm & Halby.

**References**

Boleij, M., Pabst, M., Neu, T. R., van Loosdrecht, M. C. M., & Lin, Y. (2018). Identification of Glycoproteins Isolated from Extracellular Polymeric Substances of Full-Scale Anammox Granular Sludge. *Environmental Science & Technology*, *52*(22), 13127–13135. https://doi.org/10.1021/acs.est.8b03180

Edhager, A. V., Povlsen, J. A., Løfgren, B., Bøtker, H. E., & Palmfeldt, J. (2018). Proteomics of the Rat Myocardium during Development of Type 2 Diabetes Mellitus Reveals Progressive Alterations in Major Metabolic Pathways. *Journal of Proteome Research*, *17*(7), 2521–2532. https://doi.org/10.1021/acs.jproteome.8b00276

Tahkola, K., Ahtiainen, M., Kellokumpu, I., Mecklin, J.-P., Laukkarinen, J., Laakkonen, J., Kenessey, I., Jalkanen, S., Salmi, M., & Böhm, J. (2021). Prognostic impact of CD73 expression and its relationship to PD-L1 in patients with radically treated pancreatic cancer. *Virchows Archiv*, *478*(2), 209–217. https://doi.org/10.1007/s00428-020-02888-4
